# Neural dynamics in the rodent motor cortex enables flexible control of vocal timing

**DOI:** 10.1101/2023.01.23.525252

**Authors:** Arkarup Banerjee, Feng Chen, Shaul Druckmann, Michael A. Long

**Affiliations:** NYU Neuroscience Institute, New York University Langone Health, New York, NY 10016, USA; Department of Otolaryngology, New York University Langone Health, New York, NY 10016, USA; Center for Neural Science, New York University, New York, NY 10003, USA; Cold Spring Harbor Laboratory, Cold Spring Harbor, NY 11724, USA; Department of Applied Physics, Stanford University, Stanford, CA 94305, USA; Department of Neuroscience, Stanford University, Stanford, CA 94304, USA

**Author notes:** Co-first author (Arkarup Banerjee, Feng Chen).

## Abstract

Neocortical activity is thought to mediate voluntary control over vocal production, but the underlying neural mechanisms remain unclear. In a highly vocal rodent, the Alston’s singing mouse, we investigate neural dynamics in the orofacial motor cortex (OMC), a structure critical for vocal behavior. We first describe neural activity that is modulated by component notes (approx. 100 ms), likely representing sensory feed-back. At longer timescales, however, OMC neurons exhibit diverse and often persistent premotor firing patterns that stretch or compress with song duration (approx. 10 s). Using computational modeling, we demonstrate that such temporal scaling, acting via downstream motor production circuits, can enable vocal flexibility. These results provide a framework for studying hierarchical control circuits, a common design principle across many natural and artificial systems.

## Introduction

Many species exert voluntary control over vocal production, allowing rapid flexibility in response to conspecific partners or other environmental cues [1, 2]. Neocortical activity observed across a range of species [3–8] has been proposed to be important for executive control of vocalization [9–12]. For instance, cortical neurons are preferentially active when non-human primates vocalize in response to a conditioned cue [6]. In contrast, the primary vocal motor network consisting of evolutionarily conserved brain areas in the midbrain and brainstem [10–14] is sufficient to generate species-typical sounds. Pioneering work in squirrel monkeys [15] and cats [16] as well as recent studies in laboratory rodents [17–19] have identified many such areas, including the periaqueductal grey and specific pattern generator nuclei in the reticular formation. While these subcortical vocal production mechanisms have been well-characterized, much less is known about how cortical activity contributes to vocal production and flexibility.

To address this issue, we focus our attention on the highly tractable vocalizations of a Costa Rican rodent [20]: the Alston’s singing mouse (Scotinomys teguina, **Fig. 1a**). Singing mice produce a temporally patterned sequence of notes (approx. 20 to 200 ms) that become progressively longer over many seconds (e.g., **Fig. 2a**), henceforth referred to as a song. Singing mice can flexibly adjust their song duration in response to many internal [21] and external [22] factors, including social context [20]. Recently, we discovered that a specific forebrain region, the orofacial motor cortex (OMC), is crucial for vocal behavior in this species [20]. Electrical stimulation of OMC disrupted or paused ongoing singing, and its pharmacological inactivation abolished vocal interactions and significantly reduced variability in song duration [20]. A major gap in understanding, however, concerns the nature of the cortical activity that drives this ethologically relevant vocalization. We therefore performed the first electrophysiology recordings in singing mice to assess the impact of OMC dynamics on vocal production and flexibility.

**Fig. 1.**
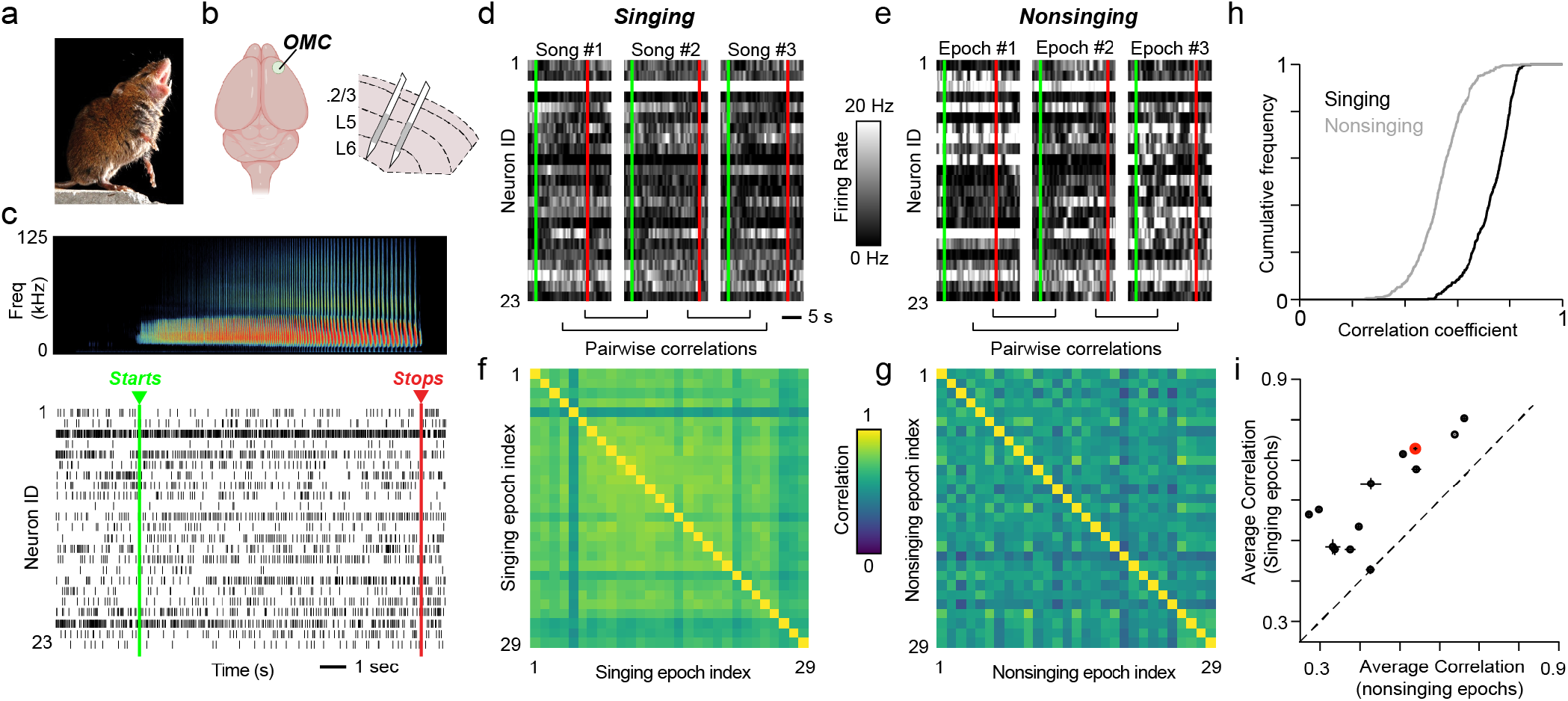
Reliable cortical population activity during singing in S. teguina. **(a)** S. teguina singing (Photo credit: Christopher Auger-Dominguez). **(b)** Schematic of S. teguina brain highlighting the recording site (i.e., orofacial motor cortex, or OMC) as well as the positioning of electrodes (gray shaded region). **(c)** Spiking activity from 23 simultaneously recorded OMC neurons during song production. The sonogram at top depicts S. teguina song. Neurons with mean firing rates less than 1 spikes/s are excluded for visualization purposes. **(d** and **e)** Firing rates of OMC neural ensemble from (c) during three singing epochs (d) compared with equally timed epochs recorded outside of song (e). For plots (c) through (e), green and red dashed lines mark the beginning and end of the song, respectively. **(f)** and **(g)** For the example session, pairwise correlations of the joint activity of the OMC ensemble recorded across all singing (f) and nonsinging (g) epochs. Dimensions of this matrix reflect the total number of songs in this session (n = 29). **(h)**Correlation values across all songs are significantly higher during singing compared with nonsinging (one-sided Welch’s t-test, p = 3.0 × 10^−139^). (i) Average correlation values for each recording session (mean ± S.E.M., n = 13 sessions, 4 mice). Red point refers to example session shown in (c)-(h).

**Fig. 2.**
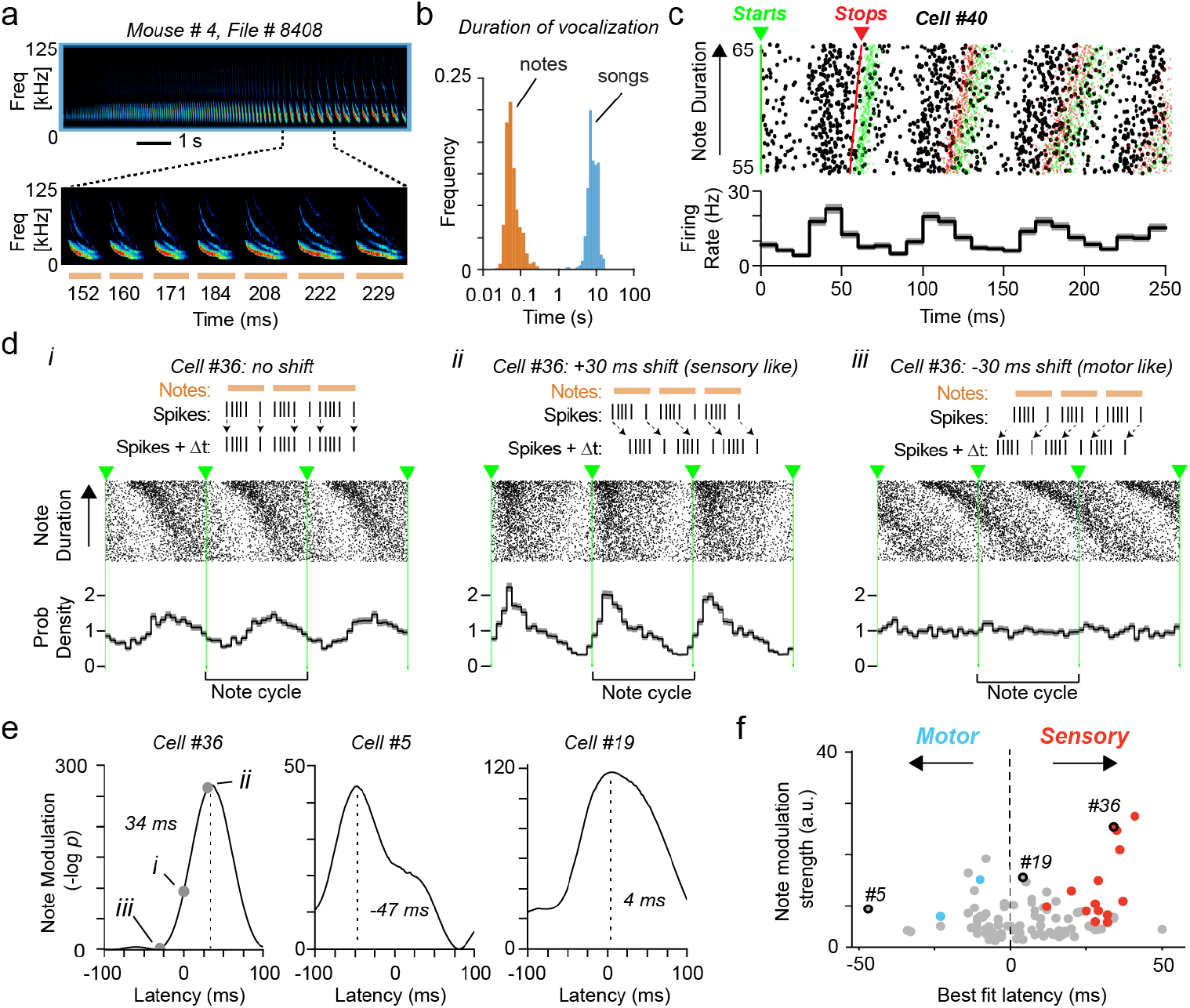
Note-related activity of OMC neurons. **(a)** At top, singing behavior in a single S. teguina example song. At bottom, an expanded view of 7 notes from the above example. Horizontal lines represent the timing of notes, and the durations for each note (in ms) are provided below. **(b)** Histogram of note (n = 30,540) and song (n = 305) durations plotted on a logarithmic axis across all recorded mice in this study (n = 4). **(c)** Spiking activity corresponding to note timing for an example neuron. For visualization, the spike raster plot was restricted to notes within a range of 55 to 65 ms (full range: 31.4 to 175.9 ms). Green and red ticks indicate the onset and offset of notes, respectively. **(d)** Spiking activity of an example neuron linearly warped to a common note duration (onsets indicated by dashed green lines). Rasters (top) and spike probability density plots (bottom) are provided for the recorded spike trains (i) and after imposing a ‘sensory’ (- 30 ms) (ii) or ‘motor’ (+ 30 ms) (iii) offset. **(e)** Modulation strength and offset values for three example neurons. Gray circles and roman numerals in plot for Cell 36 refer to the corresponding panels depicted in (d) (see Methods). **(f)** Summary plot showing the best-fit latency (restricted to ± 50 ms) corresponding to the maximum note modulation strength for 96 neurons. Gray symbols represent cases that are not significantly different from zero, and red (n = 15) and blue (n = 2) symbols represent points with sensory and motor off-sets, respectively. The three example cells depicted in (e) are indicated.

## Results

### High-density silicon probe recordings in freely behaving singing mice

We recorded OMC neural activity during vocal production in four adult male S. teguina using high-density silicon probes (Cambridge NeuroTech or Diagnostic Biochips) (**Fig. 1b, c**). Electrodes were inserted to a final depth of 600-1000 µm, such that most recording sites were in the ventral portion (i.e., motor output layers) of OMC. We used this approach to monitor neural activity continuously over 3 to 20 days, and 13 sessions with robust vocal behavior (duration: 10.4 ± 5.7 hours, mean ± SD) were analyzed further. During these recording sessions, singing mice produced songs both spontaneously (n = 226) and in response to the playback of a conspecific vocalization (n = 79). For this study, which focuses on vocal production, we combined data across these conditions, yielding a total of 23 ± 17 (mean ± SD) songs per session (range: 8 to 72). In total, we recorded from 375 neurons (29 ± 11 per session, mean ± SD) whose spiking was stably monitored throughout those recording sessions (see Methods).

### OMC spiking is modulated during vocal production

We began by examining whether OMC neural activity was related to singing behavior. Although song-related spiking patterns often differed across neurons (e.g., **Fig. 1c**), we found that the ensemble activity of simultaneously recorded OMC neurons was similar across song epochs compared to non-singing periods (**Fig. 1d, e**). Since each session consisted of multiple songs, we calculated the correlation values of OMC ensemble activity across all pairs of songs and found them to be significantly greater compared to nonsinging epochs in the example session (**Fig. 1f-h**) as well as across all recording sessions (Corr_singing_ = 0.61 ± 0.11, Corr_nonsinging_ = 0.44 ± 0.12, p = 2.76 × 10^−6^, paired t-test) (**Fig. 1i**). Taken together, we find that OMC population activity is consistently modulated during song production.

Since OMC ensemble activity displayed reliable neural dynamics during singing, we next proceeded to characterize song-related spiking in individual OMC neurons. Each song is composed of a series of notes (**Fig. 2a, b**); therefore, neural activity could a priori be related to the production of each note at a fast timescale (approx. 100 ms), or it could follow slower dynamics at timescales comparable to the entire song (approx. 10 s). By statistically comparing neural activity during vocal production (versus nonsinging epochs), we found that 29.6 percent of neurons (111 out of 375) were correlated with notes (**Extended Data Fig. 1a-c, Fig. 2c**) while 35.5 percent (133 out of 375) of neurons displayed dynamics spanning the entire song (**Extended Data Fig. 1d-f**, see Methods), and 13.1 percent were active at both timescales. Therefore, more than half of individual OMC neurons were significantly modulated with some aspect of singing behavior.

### Note-related responses of OMC neurons

Cortical activity has been shown to represent relevant kinematic features (e.g., velocity and force of effector muscles) for many movements [23]. Applying this framework to vocal production, we would expect OMC neurons to show phasic activity patterns preceding each note. To determine the relationship of OMC firing and note production, we linearly warped spiking activity to both the onset and offset of notes (**Fig. 2d**). A close inspection of note-related neurons revealed a diverse relationship between spike timing and note duration. For instance, in some cases, there appeared to be a systematic shift in the spike timing as note durations increased (e.g., **Fig. 2di**), which may arise from systematic offsets between neural activity and note production. Specifically, if this shift were due to a motor delay, or the timing needed for premotor signals to result in a behavioral change, activity would precede the production of notes [24]. Conversely, if the timing shift were due to sensory feedback, spiking activity would lag note production [25].

To explore these possibilities, we systematically varied the timing of spikes with respect to the audio recordings (**Fig. 2d, Extended Data Fig. 2a, b**) and determined the time lag that resulted in the most consistent alignment with notes (**Fig. 2e**, see Methods). Among the population of note modulated neurons, shifts resulted in significantly better alignment between neural activity and note phase in 25 cases (**Fig. 2e, f**, bootstrap p < 0.01, see Methods). Of these, 23 were consistent with sensory shifts and only 2 with motor offsets (**Fig. 2f, Extended Data Fig. 2e**). Based on the relative timing of neural activity and behavior, less than 1 percent (2 out of 375) of all recorded OMC neurons have a response profile consistent with a motor command for note production. Therefore, while we find phasic note-related activity in OMC, it is unlikely to be directly involved in the production of individual notes.

### Precise temporal scaling of OMC neural dynamics with song duration

We next explored an alternative schema based on hierarchical control in which OMC population dynamics is dominated by a set of motor primitives (i.e., distinct patterns of neural activity) which do not directly represent movement kinematics [26]. In this view, motor commands for note production are determined by downstream vocal pattern generators driven by time-varying OMC activity spanning the duration of the song, a dynamical systems framework that has been proposed in other motor control studies [27, 28]. Therefore, we broadened our view to examine the extent to which neural activity relates to the structure of the produced song at timescales comprising the entire song duration (approx. 10 s).

We tested how OMC neural dynamics covaries with song duration, which can substantially differ across renditions (**Fig. 3a**). The activity of individual neurons may evolve with identical timing regardless of song duration and therefore be correlated with ‘Absolute Time’ (**Fig. 3b**). Consequently, dynamics associated with shorter songs would simply look like truncated versions of those observed during longer songs. Alternatively, OMC neurons could reflect ‘Relative Time’ (**Fig. 3c**), in which neural activity expands and contracts to track the progression through longer and shorter songs, respectively. To test these models, we analyzed trial-to-trial differences in song duration across renditions (average variation: 139.9 percent, n = 13 sessions, e.g., **Fig. 3a**) and used a similarity analysis to compare the firing patterns of each modulated neuron after the timing of activity had been linearly warped to align the onset and offset of song (**Fig. 3d, e, Extended Data Fig. 3**). The Absolute Time model would predict a higher degree of correlation when maintaining original timing and comparing initial portions of longer songs to shorter songs, while the Relative Time model suggests the opposite (i.e., higher correlation after warping). We therefore directly compared these two scenarios and found that the explained variance of single trial firing rates was significantly greater in the warped condition compared with the unwarped condition (p = 7.5 × 10^−7^, one-sided paired t-test) (**Fig. 3f**), supporting the Relative Time model of OMC neural dynamics.

**Fig. 3.**
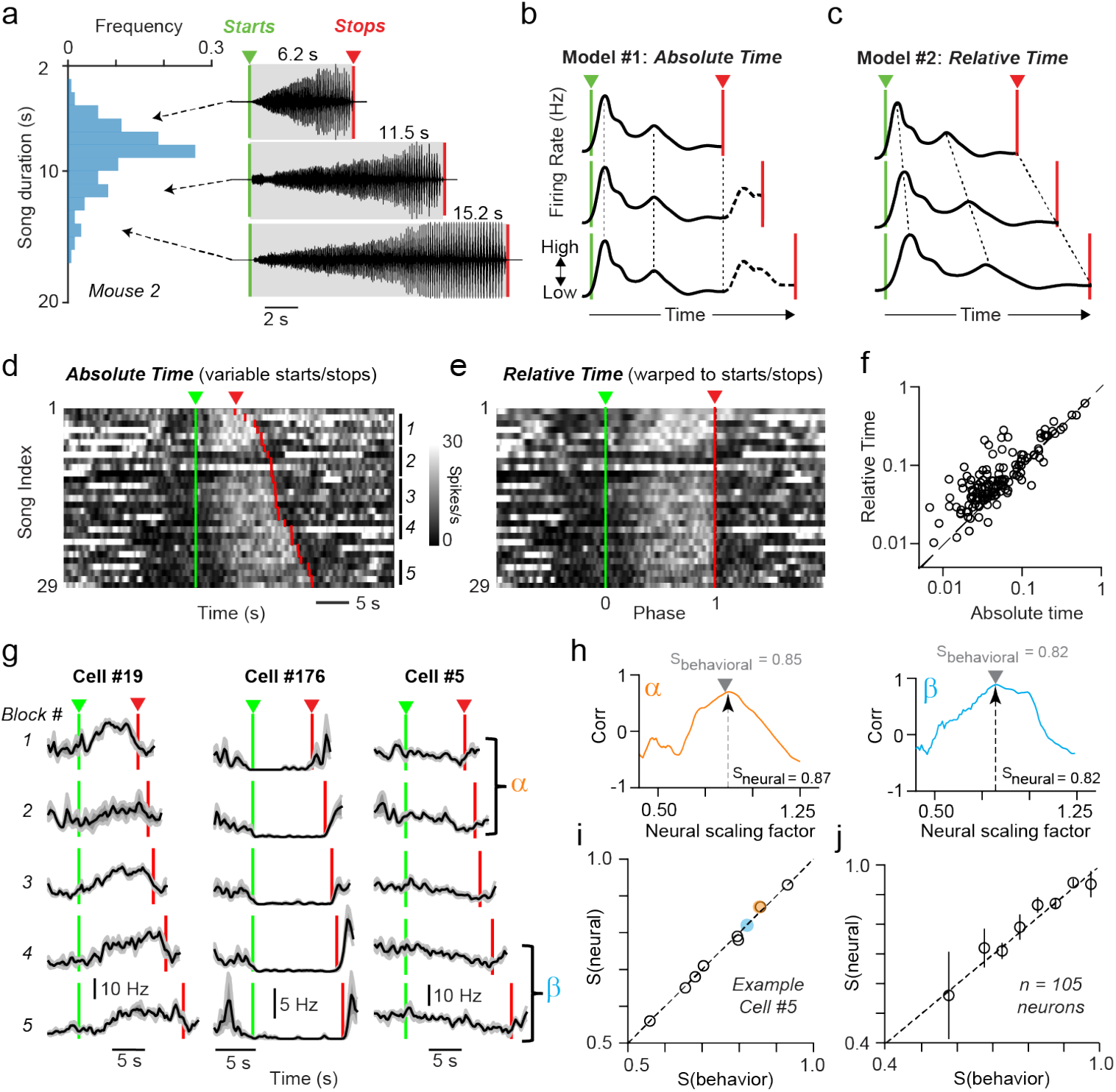
Scaling of neural activity with song duration. **(a)** Duration of all songs (n = 143) produced from one example mouse (left). Raw waveforms for three example songs of different durations (right). **(b** and **c)** Hypothetical time-varying neural activity from a single neuron as predicted by the Absolute Time (b) and Relative Time (c) models for three songs of varying durations. **(d)** and **(e)** Spiking responses for a single neuron across 29 songs aligned to the start of the song (d) or temporally warped to the beginning and end of the song (e). **(f)** Comparison of explained variance for 133 song-modulated neurons across trials using recorded song times (Model 1, x-axis) and following temporal warping (Model 2, y-axis). Data are better fit by Model 2 (one-sided paired t-test, p = 3.95 × 10^−7^). **(g)** Peri-song time histograms (PSTHs) for three example neurons. Each trace represents an average of 4-21 similarly timed trials. Cell 19 is the same neuron shown in and (e). Song blocks used to calculate consensus firing rate profiles are indicated by numbers and vertical lines. **(h)** Two example pairwise comparisons of the instantaneous firing plots from (g). For each pair, the black arrow indicates scaling factor with maximum correlation (S_neural_), and the gray arrow shows the ratio of song times (S_behavioral_). **(i)** and **(j)** All pairwise comparisons (n = 10) of S_neural_ and S_behavioral_ for the example neuron (i) (colored circles refer to panels in (h)) and for the entire population (n = 105 neurons, x-axis bin size: 0.05) **(j)** The error bars refer to the standard error of the median estimated by bootstrapping.

To further quantify the magnitude of time scaling for each neuron, we generated a consensus neural activity profile for songs with similar durations (**Fig. 3g, Extended Data Fig. 3a-c**, see Methods). For each pair of blocks, we compared the neural activity profiles to determine the scaling factor that maximized the pairwise correlation (e.g., **Fig. 3h**), which we call the neural scaling factor (S_neural_). If the optimal neural scaling (i.e., the ratio of activity profiles leading to the highest correlation value) matched the relative ratio of associated song durations (S_behavioral_), then the S_neural_/S_behavioral_ slope is expected to be 1 (equivalent to the Relative Time model). When S_neural_ was plotted against the behavioral scaling factor (i.e., ratio of the associated song durations, S_behavioral_), we found them to be linearly proportional (**Fig. 3i, j**). Across all the neurons, the neural scaling/behavioral slope was 1.01 ± 0.01 (n = 659 pairs, 105 neurons, **Fig. 3j**, see Methods). For comparison, the Absolute Time model would predict a slope of 0. This result demonstrates that activity of individual OMC neurons linearly stretches or compresses by a magnitude determined by the ratio of the song durations, enabling OMC activity to precisely track the proportion of elapsed song.

### Diverse individual neuron dynamics in OMC

What are the motor primitives observed in OMC during vocalization? Since OMC circuit activity precisely scales with song duration, we linearly warped the firing rates of song-modulated neurons to both the onset and offset of song. Using this strategy, we observed diverse firing patterns within the OMC during vocalization (**Fig. 4**). To quantify this heterogeneity, we performed hierarchical clustering (**Fig. 4a**, see Methods) and found that 28.6 percent of neurons increased firing during song production while the remainder were suppressed. Further analyses of their response profiles revealed 8 distinct clusters of neurons (**Fig. 4a, b**). We observed that some neurons exhibited transient responses coincident with song onset (Cluster 7), song offset (Cluster 6), or both (Cluster 2), and other neurons showed more persistent increases (Clusters 1, 3) or decreases (Clusters 4, 5, 8) in neural activity during singing. Overall, neurons were responsive throughout the duration of the song and not just at song initiation and termination, consistent with moment-by-moment control of ongoing song production. We conclude that the population of OMC neurons that keep track of Relative Time (i.e., phase) shows diverse firing patterns during song production.

**Fig. 4.**
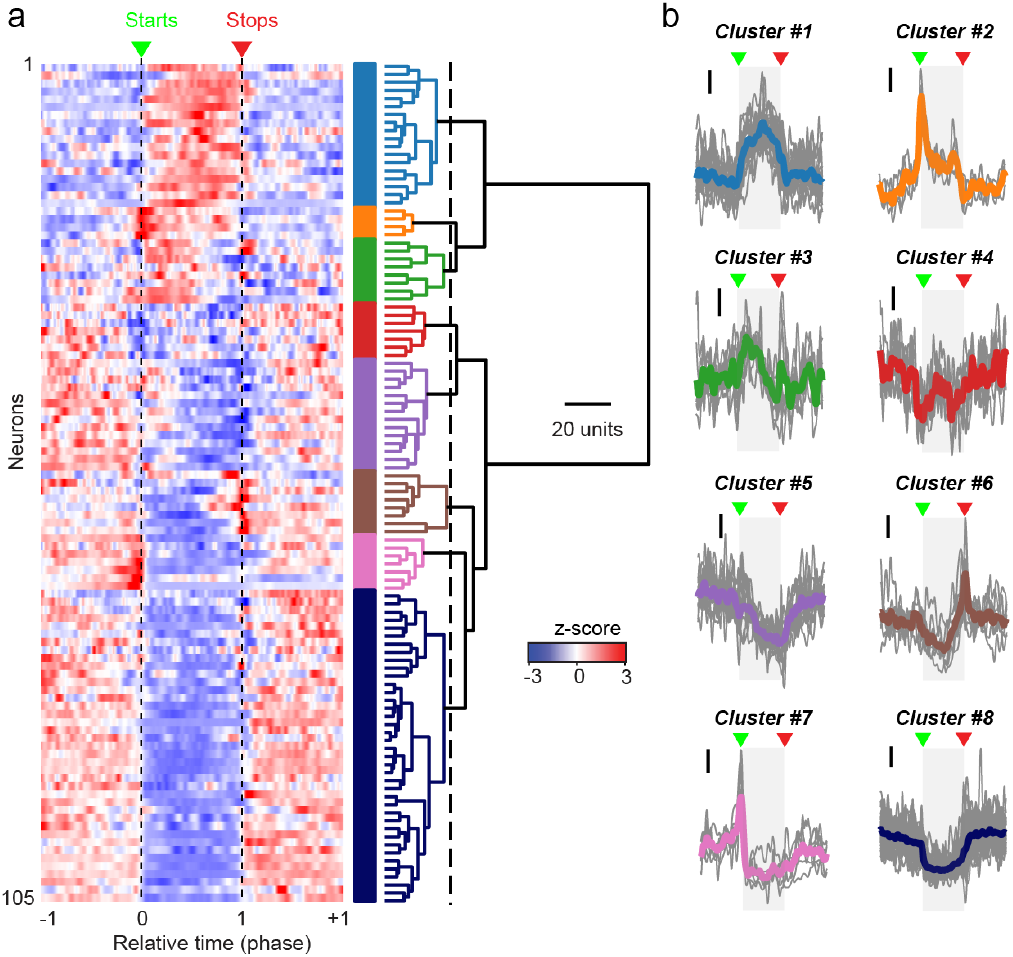
Diverse categories of OMC firing patterns during singing. **(a)** A hierarchical clustering plot describing the response profiles of OMC neurons whose activity was modulated during singing (see Methods, n = 105 neurons). Individual clusters are indicated by colored bars on the right. **(b)** Spiking responses for each cluster displayed as average firing rate plots. The mean activity profile of each neuron is represented with gray lines, and colored lines are average waveforms for each cluster corresponding to categories from (a). Black vertical bars indicate a normalized firing rate (z-score) of 1. Gray shaded blocks denote song epochs, with green and red arrows marking song starts and stops respectively.

### Computational model of vocal motor control

To understand how motor commands for note timing can be generated from the motor primitives described above (**Fig. 4**), we next constructed a data-driven hierarchical model that makes experimentally testable behavioral predictions. In this model, OMC does not determine note timing directly (consistent with a lack of ‘premotor’ timing in **Fig. 2**), but vocal motor control is instead shared by cortical and down-stream circuits. Inspired by our data, we posit that cortex dictates the moment-by-moment song phase and overall duration (**Fig. 3**), while the motor command for individual notes is generated by midbrain/brainstem areas comprising the primary vocal motor network (**Fig. 5a, Extended Data Fig. 4**). In the model, OMC activity provides descending synaptic drive, which influences the rate of note production in the subcortical song pattern generator (**Fig. 5b**). To account for the decreasing rate of note production with time, the synaptic drive onto the downstream note pattern generator may decrease throughout the song. We accomplish this in our model through linear weighting of OMC activity profiles directly measured in our recordings (**Extended Data Fig. 4a**) which sum up to produce synaptic drives with varying slopes (**Fig. 5b**). We model the workings of the note pattern generator such that individual notes are produced upon reaching a fixed firing rate threshold (see Methods), akin to an integrate-and-fire module. Appropriate time-scaling of cortical activity will thus result in songs of different durations without the need for modifying the note-generating mechanism (**Fig. 5b**). Importantly, this role of OMC is robust to the choice of the precise means by which note generation is implemented in the note pattern generator, either via postsynaptic adaptation mechanisms or synaptic drive from another brain region (**Extended Data Fig. 4b, c**).

**Fig. 5.**
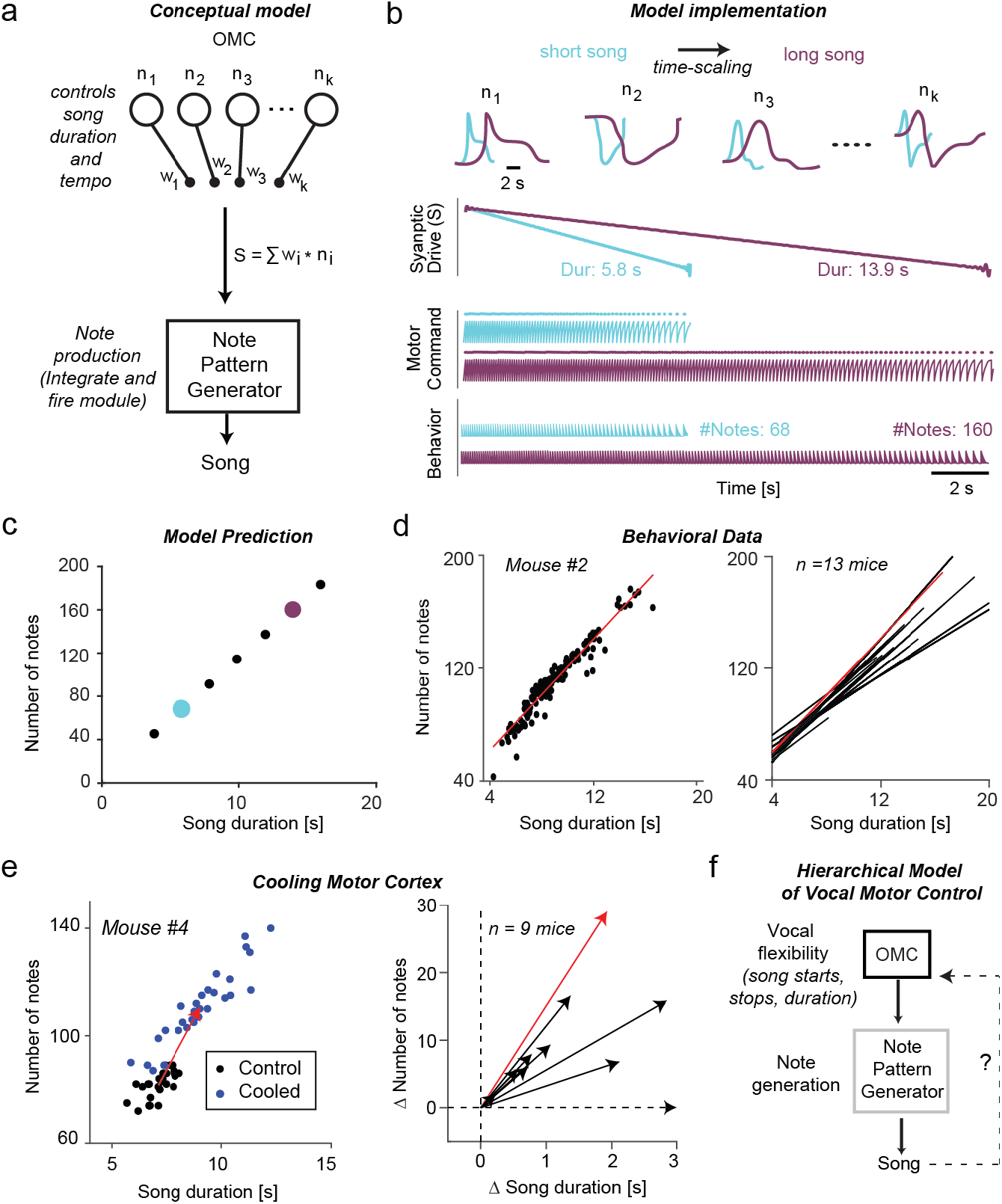
Hierarchical model of vocal motor control. **(a)** Schematic depicting shared control of vocal production, where OMC controls song duration and rate of progression while individual notes are produced by a downstream note pattern generator. The synaptic drive to the note pattern generator is derived from OMC neural activity (see **Extended Data Fig. 4**). **(b)** Activity profiles of four model OMC neurons for a long song (purple) compared to a short song (cyan). Linear summation of neural activity creates the synaptic drive to the note pattern generator. The note pattern generator is modeled as an integrate-and-fire module, such that the rate of note production depends upon the strength of the OMC synaptic input. **(c)** Model output using seven different values of time-scaling, leading to a prediction in which the number of notes linearly covaries with song duration. Cyan and purple indicate examples from (b). **(d)** The number of notes scales with song duration in an example mouse (n = 144 songs, left) as well as across the population (n = 13 mice, right). Diagonal lines at right represent linear regression fits for each individual animal. Red line indicates data from the example at left. **(e)** Cooling OMC results in a shift in both song duration and number of notes in one example animal (Mouse 4, left). The average change in song duration and number of notes as the result of cooling for each animal (n = 9 mice, right). Across all animals, OMC cooling significantly increased average song durations (control: 8.0 ± 0.3 s, cooled: 9.3 ± 0.4 s, p = 0.002, paired t-test) as well as average number of notes (control: 92.8 ± 3.2, cooled: 103.9 ± 3.5, p = 0.004, paired t-test). Red line indicates data from Mouse 4 (left). **(f)** Hierarchical model of vocal motor control, wherein OMC confers flexibility to a downstream song pattern generator.

We next test a specific behavioral prediction of our hierarchical model to assess its validity. Our model predicts that songs become longer by incorporating more notes and not by increasing the duration of individual notes (**Fig. 5b, c**). Alternately, if note timing were directly triggered by note-modulated OMC activity (**Fig. 2**), longer songs would have the same number of notes with their durations proportionately stretched, as observed in the songbird [29, 30]. We tested these predictions by examining the structure of songs produced with different durations and found that the number of notes systematically increased as a function of song duration (n = 13 animals, 4 from this study and an additional 9 from a published data set [20]) (**Fig. 5d**), a finding that strongly agrees with our hierarchical model.

We considered a directed circuit perturbation to assess whether the relationship between notes and song duration relies upon activity within OMC. We reanalyzed a data set in which OMC was focally cooled in 9 mice [20]. Previous experimental [29, 31-33] and theoretical [34] work predicts that mild focal cooling should dilate the temporal profile of OMC neural activity thereby slowing the progression of sub-cortically controlled note production. For each animal, OMC cooling resulted in an increase in both song duration (control: 8.0 ± 0.3 s, cooled: 9.3 ± 0.4 s, p = 0.002, paired t-test) as well as the number of notes (control: 92.8 ± 3.2, cooled: 103.9 ± 3.5, p = 0.004, paired t-test) (**Fig. 5e**). Therefore, OMC-cooled songs became longer by incorporating more notes, further supporting the role of OMC activity in our hierarchical model. In sum, these results suggest that cortical activity can generate the necessary vocal motor commands to account for natural variability in behavior.

## Discussion

In this study, we observed robust modulation of motor cortical activity during vocalization corresponding to two behaviorally relevant timescales: (1) phasic responses during note production (approx. 100 ms) and (2) persistent song-related dynamics (approx. 10 sec). We found that many neurons modulated at the faster timescale exhibited a delay between note timing and spiking that could represent either sensory feedback or efference copy signals (**Fig. 5f**). Sensory feedback is known to be important in animal and human vocal motor control [35–38], and a systematic perturbation of sensory streams (e.g., auditory, proprioceptive) [39] could test whether these signals are important in similar control processes in the singing mouse. Nevertheless, our time-shift analysis, modeling, and perturbation results confirm that these fast-varying responses in OMC do not reflect vocal motor commands to produce individual notes. At the slow timescale, responses were heterogeneous (e.g., transient at song onsets, ramping responses, etc.) and appear to reflect a set of motor primitives related to the control of song duration and the rate of note production. Future work will determine whether these spiking profiles map onto specific neuronal cell types in the OMC defined by critical circuit features, such as their output targets, as seen in motor cortical circuits in the laboratory mouse [40–42].

These results provide a striking example of how motor cortical dynamics can modulate song production, perhaps reflecting a voluntary mechanism of generating adaptive vocal flexibility. To accomplish this moment-to-moment control, our cortical recordings support a model in which OMC acts hierarchically via downstream song pattern-generator circuits (**Fig. 5f, Extended Data Fig. 4b, c**), likely corresponding to regions that have been recently characterized in the laboratory mouse [17–19] and appear to be highly conserved across vocalizing species [10, 11]. The hierarchical model proposed here is consistent with our previous work, where we found that OMC inactivation did not abolish singing but significantly reduced the variability in song durations [20], suggesting that activity in OMC is providing necessary input to the brainstem to generate socially appropriate vocalizations (**Extended Data Fig. 4b, c**). Future work is needed to determine the full song circuit in the singing mouse and elucidate the synaptic mechanisms by which OMC influences downstream vocal production circuits.

The singing mouse vocal control network appears to operate in a partially autonomous hierarchical configuration – a successful design principle for biological and artificial systems – wherein a higher order modulator (i.e., OMC) extends the capabilities of lower-level motor controllers (i.e., note production circuitry) without being necessary for generating the basic motor program [43–45]. Such an arrangement enables behavioral flexibility without relying upon synaptic plasticity in downstream motor patterning circuits. Similar mechanisms have been observed when animals are trained to keep track of time [46–51] or in primate cortex during cycling tasks at different speeds [52]. Our results extend the scope of this temporal scaling algorithm over an expanded time window (approx. 10 s) and to a new domain: controlling vocal flexibility in mammals. Despite its ubiquity, the neural mechanisms contributing to temporal scaling are not well-understood, though several ideas have been proposed, including feedback loops [46, 51] and neuromodulatory gain control [53]. The OMC circuit in the singing mouse offers a valuable opportunity to examine these and other circuit features for generating motor flexibility in the context of an ethologically-relevant behavior.

## Acknowledgements

We thank Steve Shea, Florin Albeanu, Walter Bast, Joseph del Rosario, Hadas Sloin, and members of the Long and Banerjee laboratories for comments on earlier versions of this manuscript. Abby Paulson provided technical assistance.

## Funding

National Institutes of Health grant R01 NS113071 (MAL, SD)

Simons Collaboration on the Global Brain (MAL, SD) Searle Scholars Program (AB)

Simons Foundation Junior Fellows Program (AB)

## Author contribution

Conceptualization: AB, MAL

Methodology: AB, FC, SD, MAL

Investigation: AB, MAL

Visualization: AB, FC, SD, MAL

Funding acquisition: AB, SD, MAL

Project administration: SD, MAL

Supervision: SD, MAL

Writing – original draft: AB, MAL

Writing – review and editing: AB, FC, SD, MAL

## Data and materials availability

Further information and requests for resources and reagents should be directed to and will be fulfilled by the Lead Contact, Michael Long. This study did not generate new unique reagents. The data sets generated during this study are available upon request from the Lead Contact.

## Supplementary Materials

## MATERIALS AND METHODS

### Animals

All procedures were conducted in accordance with protocols approved by the Institutional Animal Care and Use Committee of NYU Langone Medical Center. Animals used in the study were adult (> 3 months) male laboratory-reared offspring of wild-captured *Scotinomys teguina* from La Carpintera and San Gerardo de Dota, Costa Rica. Mice were maintained at 22 ± 3 °C with a 12:12 L:D cycle.

### Behavioral recordings

*S. teguina* were housed in individual recording chambers (Med Associates) lined with sound insulation foam (Soundproof Cow). Vocalizations were recorded using a condenser microphone (Avisoft Bioacoustics CM16/CMPA) placed within home cages. Acoustic signals were sampled at 250 kHz and digitized with Avisoft UltraSoundGate 116Hb. For playback experiments, we used an ultrasonic tweeter (Vifa), as described previously [1]. To precisely align the audio and electrophysiology signals, each data stream was additionally recorded continuously into an INTAN recording system at a fixed sampling rate between 20-30 KHz.

### Silicon-probe recordings

Chronic recordings were performed using either 64-channel (Cambridge Neurotech, E-1) or integrated 128-channel high-density silicon probes (Diagnostic Biochips, 128-5). Prior to surgery, probes were mounted to a plastic microdrive (NeuroNexus, d-XL) and a stainless-steel ground wire (0.001”, A-M systems) was soldered to the reference of the headstage, which was held in place by a custom-made 3D printed enclosure (Formlabs). For all surgical procedures, mice were anesthetized with 1-2% isoflurane in oxygen and placed in a stereotaxic apparatus. Neural activity of freely moving singing mice was recorded using an electrically assisted commutator (Doric Lenses) and the RHD USB Interface Board or RHD Recording Controller (Intan Technologies). For all chronic recordings, silicon probes were implanted directly into the OMC using the following stereotaxic coordinates: +2.25 mm anterior to bregma, +2.25 mm lateral to the midline. This location represents the center of the OMC region identified by electrical microstimulation [1]. The ground wire was inserted between the skull and the dura above the visual cortex or cerebellum contralateral to the probe implantation. Silicon elastomer (Kwik-Cast, WPI) was applied to the craniotomy once the probe was inserted to the desired depth (1 mm for OMC). The microdrive and the enclosure were secured to the skull with dental acrylic and Metabond cement (Parkell). Animals were monitored and allowed to recover for three to seven days prior to the start of electrophysiology experiments. Spike detection and clustering were performed using KiloSort software [2] and manual post-processing (merging/ splitting of clusters) was performed using phy [3]. Clusters that drifted during the recording session were not included in further analyses. Spike times of all clusters were aligned to onsets and offsets of individual notes or songs as specified below.

### Behavioral annotation of acoustic parameters

We analyzed song structure using custom software (MATLAB) as described previously [1]. Briefly, we first smoothed the sound waveform with a 4-ms sliding window. We then identified individual notes, which typically exhibited an absolute intensity threshold corresponding to 25-40 dB below the mouse’s loudest note. Exact note start times and stop times were calculated based on the maximum intensity of each note, such that onsets and offsets were first and last crossings of 1% (20 dB quieter) of each note’s maximum intensity. Note duration was calculated as the difference between the offset and the onset for each note. Song duration was defined as the difference between the offset of the last note and the beginning of the first note. For each song, the number of notes was plotted against the overall song duration. For each animal, linear regression (MATLAB function: polyfit) was used to describe how the number of notes vary as a function of song duration. For reanalysis of the previously published cooling data set [1], the number of notes for each song was plotted against the song duration for both control and cooled conditions. A small minority of songs shorter than five seconds, which often had breaks, were ignored. To summarize the effect of cooling, for each animal, the difference between the average number of notes before and after cooling was plotted against the difference of song durations before and after cooling. Since the average song duration varies for individual animals, the difference between cooled and control conditions (∆ notes and ∆ song duration) were plotted as opposed to the absolute values.

### Correlation analysis of neuronal ensembles during singing

We performed a correlation analysis for each session individually. We estimated the firing rates from the spike trains using a Gaussian kernel (σ = 0.2 s). For correlation analyses, we chose the window size based on the longest song duration *T*_max_ in that session. To better capture the modulation at the onset and offset, we included 2 s before the song onset and 2 s after the song offset, so the total window size is *T*_max_ + 4 *s*. In this time window, for each song in the session, we sampled every 200 ms from the estimated firing rates to construct the peri-song time histograms (PSTHs). We concatenated the PSTHs from all the neurons for each song into a single vector. The correlation matrix was then constructed by taking the correlation between all pairs of songs. For nonsinging epochs, we performed the same analysis but with song timing (onsets) replaced by control epochs set to be 30 seconds after the song offsets. For each session we averaged the off-diagonal elements in the correlation matrix and performed a one-sided paired t-test to determine the significance.

### Selection of note and song modulated neurons

#### Note modulated neurons

Within a song, consecutive notes usually have short gaps between them (~1/3 of note duration, e.g., **Fig. 2a**). We define a note cycle (*T*_cycle_) as the time between the subsequent note onsets. Some songs may have short pauses. To distinguish actual note cycles from pauses, we required the note cycle duration to be less than 3 times the note onset-offset duration. All the analyses on notes shown in this paper were performed with note cycles that passed the above criterion. We verified that our results are robust if we change note cycles to be the time from note onset to offset or the time between the offsets of successive notes. Because notes have variable durations, our analyses were carried out after warping spiking activity to align onsets and offsets, which enabled the calculation of phase tuning. We defined note phase as the relative time within a note cycle, 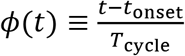. To select note-modulated neurons, we summarized the spike phases for all the notes and used the Rayleigh *z*-test (*α* = 0.01) to test against the null hypothesis that the spikes within each note cycle were uniformly distributed.

#### Song modulated neurons

We selected the song-modulated neurons initially without warping, i.e., in Absolute Time. Because each song within a session has a different duration, and the different durations could affect estimations of variance, we used the same window size for all songs. Specifically, we chose the window size based on the shortest song duration *T*_min_ in that session. We performed statistical tests twice: once for song onset alignment and once for song offset alignment (**Extended Data Fig. 1d**). To better capture the modulations at song onsets or offsets, we include 2 s before the song onsets or 2s after the song offsets. For song onset alignment, we calculated the averaged firing rates within the time window by counting the spikes between 2 s before the song onsets and *T*_min_ after the song onsets. For song offset alignment, we calculated the averaged firing rates within the time window by counting the spikes between *T*_min_ before the song offsets and 2 s after the song offsets. As a control, we created a baseline nonsinging epoch for each song by counting the spikes from 10 s to 70 s after the song offset. In rare cases when another song appeared in this time window, we excluded the song period and extended the time window to include a total of 60 s of baseline activity. We then performed a two-sided paired *t*-test (*α* = 0.01) to test the null hypothesis that the firing rates within a song were the same as baseline firing rates.

### Analysis of note-related neural activity

We found that for many neurons the time course of modulation by notes had a peak that shifted with note duration (e.g., **Fig. 2di**). One possible explanation is that there exists a latency in absolute time between the behavioral recordings and neural activity. To quantify this offset, we reasoned that the optimal latency should give the strongest modulation that is characterized by the *p*-value of the Rayleigh *z*-test. We defined the modified phase as 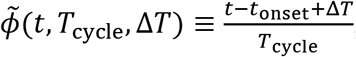, where Δ*T* is the fixed latency in absolute time, *T*_cycle_ is the note cycle duration, and *t*_onset_ is the note onset. We performed the Rayleigh *z*-test in this modified phase frame and obtained the *p*-value as a function of the latency Δ*T*. The optimal latency was determined from 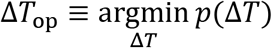. To calculate the modulation strength, we first defined the modulation vector as 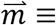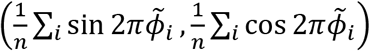, where *n* is the total number of spikes in all the note cycles. We estimated the standard error of the *L*_2_ norm of the modulation vector and denoted it as 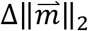. The modulation strength is then defined as 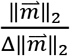.

To check whether the latency was sensory- or motor-like, we first selected neurons that had a latency that significantly differed from 0 based on bootstrapping; we randomly sampled the note cycles 1000 times to get the distribution for inferred optimal latency. We then selected neurons which had an optimal latency distribution significantly different from 0 (two sides, *α* = 0.01). We define song modulation strength by the larger absolute *t*-value of the two *t*-tests (i.e., performed on onset- and offset-aligned data).

### Analysis of song-related neural activity

To differentiate between the absolute time and relative time models, we constructed a mean template and compared the variance explained by each model. We estimated the firing rates from the spike trains using a Gaussian kernel (σ = 0.2 s) and denoted this continuous function as *r*_σ_(*t*). For the absolute time model, we set the time window to be the shortest song duration *T*min in that session and sampled every 200 ms in this window from *r*_σ_(*t*) to construct the PSTHs. This gave a matrix ***R***^abs^, which is of size (*n*_song_, 5 ∗ *T*_min_). For each neuron, the mean template was then constructed by taking averages across the rows (i.e., song dimension). We then computed the explained variance (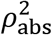 of the PSTHs about the mean template. For the relative time model, we sampled the same number (5*T*_min_) of points evenly from the firing rate function *r*_σ_(*t*) of each neuron after linear warping of time between song onset and song offset. Explicitly stated, 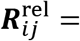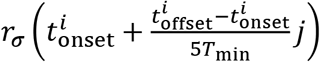, where 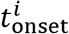 and 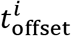 denote the onset and offset for the i^th^ song in the session. Following this, identical to above, we computed the mean template and the explained variance using ***R***^rel^ in place of ***R***^abs^.

To further quantify the degree of stretching and compression in the relative time model, we performed the following scaling analysis. For each session, we first grouped songs of similar durations using the Jenks Natural Breaks method [4]. We required a valid cluster to have at least four songs and chose the number of clusters to maximize the total number of valid clusters in each session. We then averaged the neural firing rates within each song cluster, 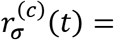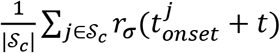, where the superscript (*c*) denotes the cluster, and *δ_c_* denotes the set of song indices in cluster c. For any two clusters (e.g., *c*_1_ and *c*_2_), the goal was to find the scaling factor *s_neural_* that gave the largest correlation between the two cluster-averaged firing rates 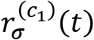 and 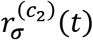. Formally, for a given scaling factor *s*, we first chose the window size 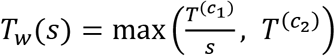, where 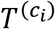 is the average song duration in that cluster. We then computed the correlation between the two cluster-averaged firing rates along the time dimension using 31 sampling points from 0 to *T*_*w*_(*s*). The optimal neural scaling is defined by 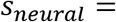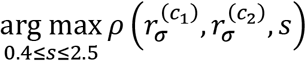. We get the behavior scaling factor from 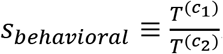. If the neural firing rates can be explained by relative time, we would get *s_neural_* ≈ *s_behavioral_*. Depending on whether 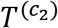 is longer or shorter than 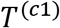, the behavior scaling factor *s_behavioral_* would be either larger or smaller than 1. To eliminate the ambiguity of these two choices of orders, we required *s_behavioral_* ≤ 1, i.e., we chose the order such that 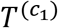 is smaller than 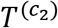. For a single neuron, we performed the scaling analysis on all possible combinations of the cluster pairs. To perform this analysis, two valid clusters per session were required (12/13 sessions met this criterion). Scaling analyses were only performed on song modulated neurons whose firing rates exceeded 1 Hz either within the song or within the control. To summarize the results, we binned *s_behavioral_* (bin size = 0.05) and plotted the median of *s_neural_* within each bin. The best fit line was estimated using quantile regression without intercept.

### Hierarchical Clustering

We estimated firing rates from spike trains using a Gaussian kernel (σ = 0.2 s) and denoted this continuous function as *r*_σ_(*t*). For the song-modulated neurons, we linearly warped their absolute time firing rates to the relative time firing rates and take the mean across songs, 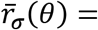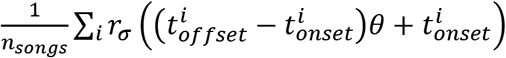. We then transformed 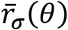 to its *z*-score. For each neuron, we sampled *z*(*θ*) from −0.2 to 1.2 with an interval of 0.01, which composes the vector representation of the neural modulation with the song. Agglomerative clustering was carried out on those vector representations. We used Euclidean distance as the affinity function. We chose the distance threshold to be 25. An average template was computed for each cluster by averaging across the neurons within the cluster.

### Computational Model

We constructed a two-step model for hierarchical vocal motor control in the singing mouse. We assumed that a note pattern generator integrates synaptic input and fires upon reaching a fixed threshold using the leaky integrate-and-fire equation:

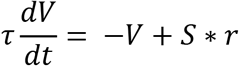

where *V* = Instantaneous voltage of the note pattern generator and *S* = synaptic drive onto the pattern generator. *r* and *τ* are the membrane resistance and the membrane time-constant respectively, with units chosen appropriately. *V* was initialized and reset to 0 mV whenever it reached a particular threshold voltage (V_th_ = 50 mV). This constituted the motor command for producing each note.

Since the rate of note production per unit time steadily decreases as the song progresses, the overall synaptic drive was required to have a negative slope. In the simplest version of the model, we assumed that the synaptic drive is entirely derived from OMC population activity. The synaptic drive was estimated using a linear combination of synaptic weights from the empirical neural data. The synaptic weights were calculated for one standard song duration (~8 s) close to the average song duration in this species. Notice that the shape of the synaptic drive (sloping down) does not require individual OMC neurons to do so. This should be interpreted as the effective influence of OMC on the note pattern generator. To generate songs of different durations (e.g., T = 4 to 16 s), OMC neural activity was time-scaled by the exact ratio of the song durations (i.e., T/8) based on our empirical result without modifying the synaptic weights. This generates steeper slopes for songs shorter than 8 s and shallower slopes for songs longer than 8 s. This model predicts that the total number of notes corresponding to each song duration increases linearly, which is recapitulated by the behavioral and cooling data. We find that this key result holds for large ranges of the values of the model parameters (*V*, *V_th_*, *S*, *r*, *τ*)

Currently, mechanistic details of the pattern generator circuit are unknown. Thus, we explore an alternative scenario by relaxing the assumption that the synaptic drive is entirely driven by OMC without any loss of generality. Its origin can be either entirely driven by OMC, or a combination of OMC and other brain areas. Moreover, the downward sloping synaptic drive can in practice result from a combination of a time-scaled duration signal and spike-frequency adaption (**Extended Data Fig. 4**).

## EXTENDED DATA FIGURES

**Extended Data Fig. 1.**
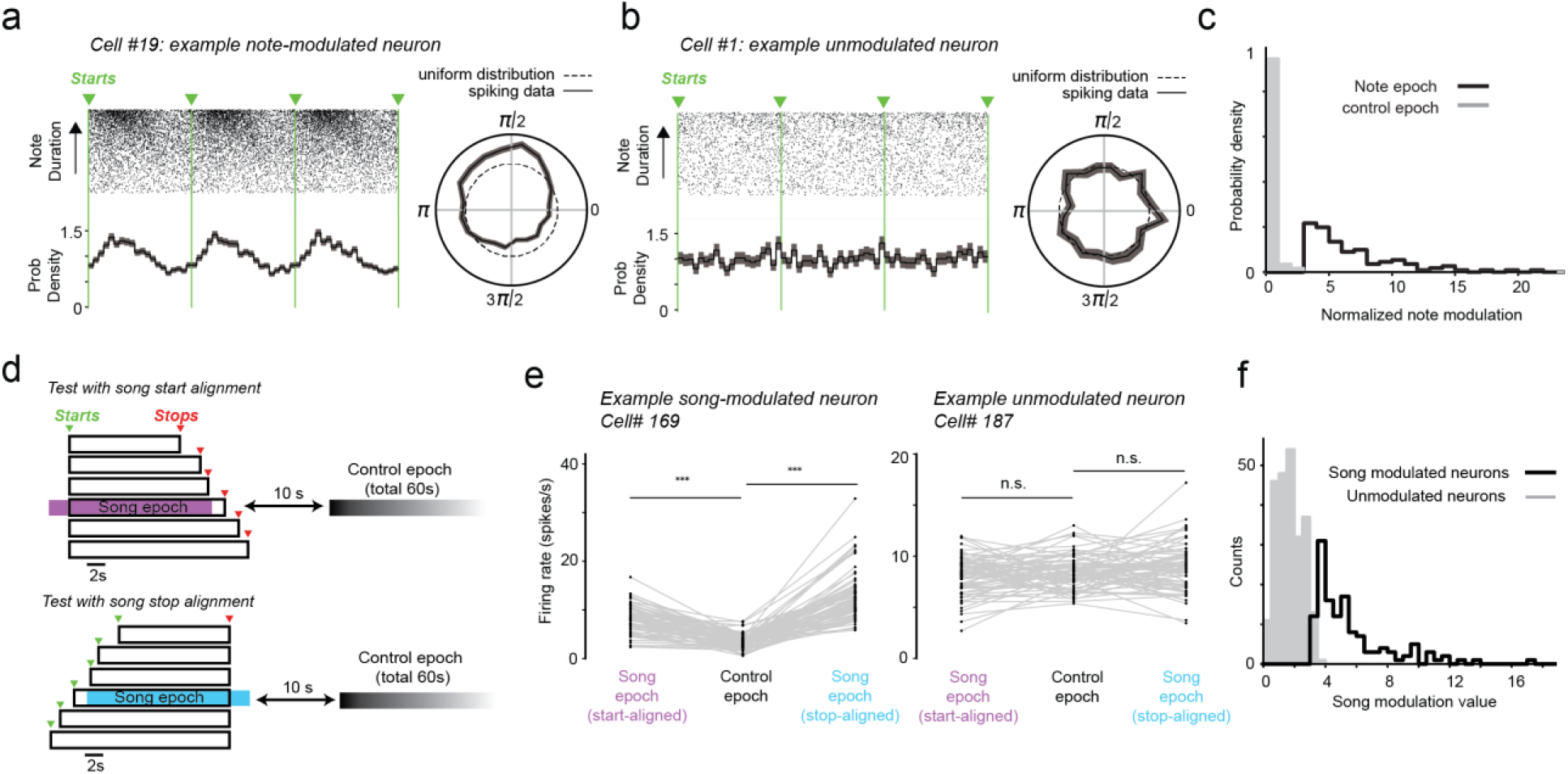
Determination of significant note- and song-related responses. (**a** and **b**) Example neurons with (a, Cell #19) and without (b, Cell #1) significant note modulation. Rasters (top) and spike probability density plots (bottom) for example neurons whose activity profiles have been linearly warped to a common note duration (onsets indicated by green lines). At right, polar plots describing the tuning of spike times with respect to the relative phase of note production. Dashed lines indicate a uniform distribution. (**c**) Histogram of note modulation (see Methods) for significantly note-modulated neurons (n = 111) compared with the same analysis applied to nonsinging epochs. (**d**) Song modulation analysis protocol. Neural activity for songs (black rectangles) are aligned either to their starts (top) or stops (bottom). The evaluation window (song epoch) begins and ends two seconds before and after the shortest song duration of that session. (**e**) The relative firing rate difference between the song-aligned spiking activity and a nonsinging period for a modulated (left, Cell #169) and unmodulated (right, Cell #187) neuron. 72 song trials are represented by separate lines for each neuron. Significance determined by bootstrap resampling (***: p < 0.01, n.s.: not significant). (**f**) Histogram of song modulation values (see Methods) for all song modulated neurons (n = 133) and those not modulated by song (n = 242).

**Extended Data Fig. 2.**
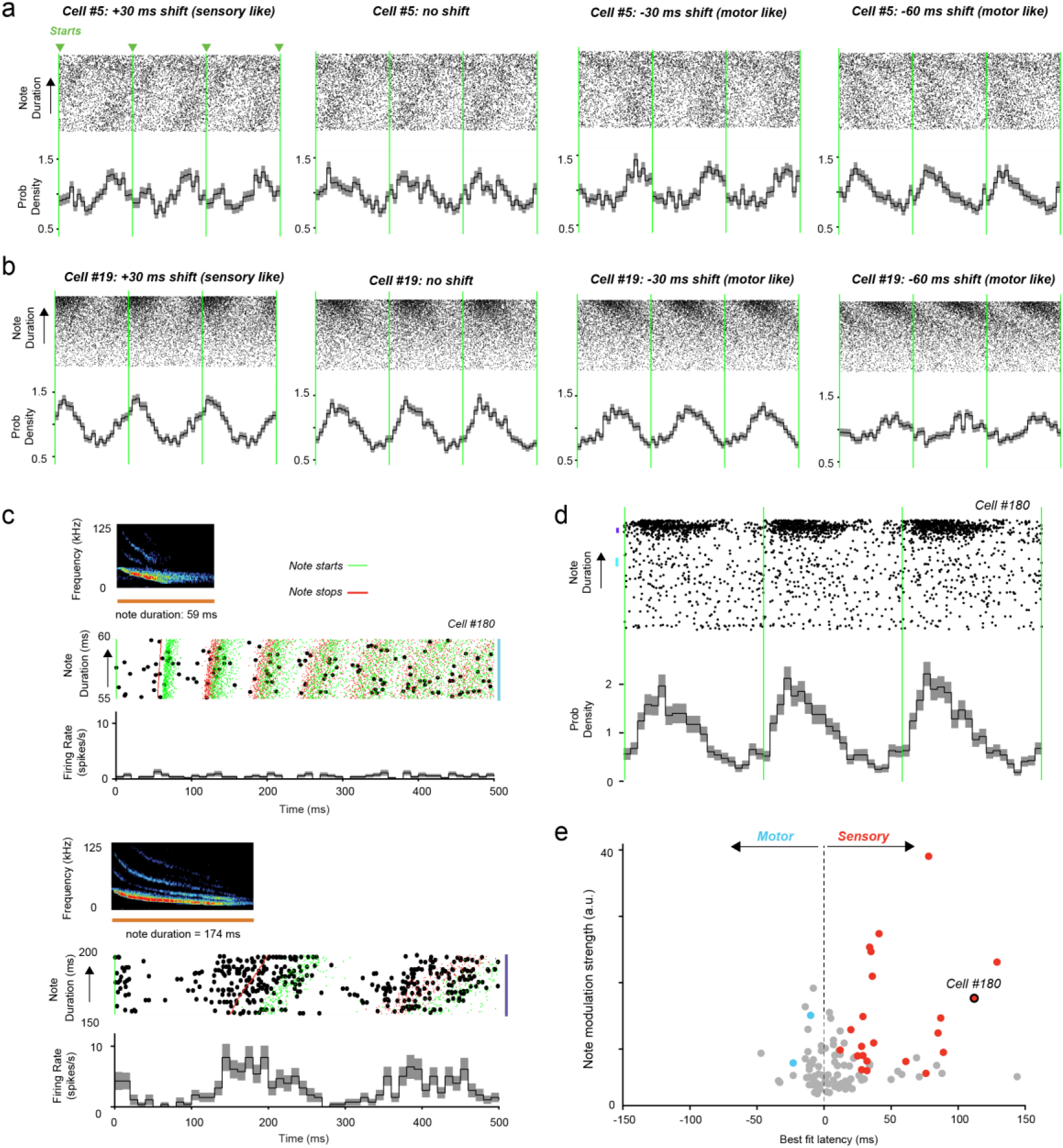
Further characterization of note-related responses. (**a** and **b**) Spiking activity of two example neurons – Cell #5 (a) and Cell #19 (b) - linearly warped to a common note duration (onsets indicated by dashed lines). At right, the alignment of spikes under normal conditions and after imposing a ‘sensory’ (− 30 ms) or two different ‘motor’ (+ 30 ms) and (+ 60 ms) offsets. Examples in (a) and (b) relate to analyses in **Fig. 2e**. (**c**) Spiking activity corresponding to note timing for an example neuron (Cell #180 from Mouse #4). For visualization, analysis was restricted to notes of prespecified durations (top: 55 to 60 ms; bottom: 150 to 200 ms, sample note sonograms provided for each range). For long note durations, robust spiking emerges near the end of each note. Green and red ticks indicate the onset and offset of notes, respectively. (**d**) Spiking activity from Cell #180 linearly warped to a common note duration (onsets indicated by dashed lines). Timing shifted by a best fit latency of 110 ms (sensory-like shift). (**e**) Summary plot (extension from **Fig. 2f**) showing the latency resulting in the maximum note modulation strength for all note modulated neurons (n = 111). Gray symbols represent cases that are not significantly different from zero, and red (n = 23) and blue (n = 2) symbols represent points with sensory and motor offsets, respectively.

**Extended Data Fig. 3.**
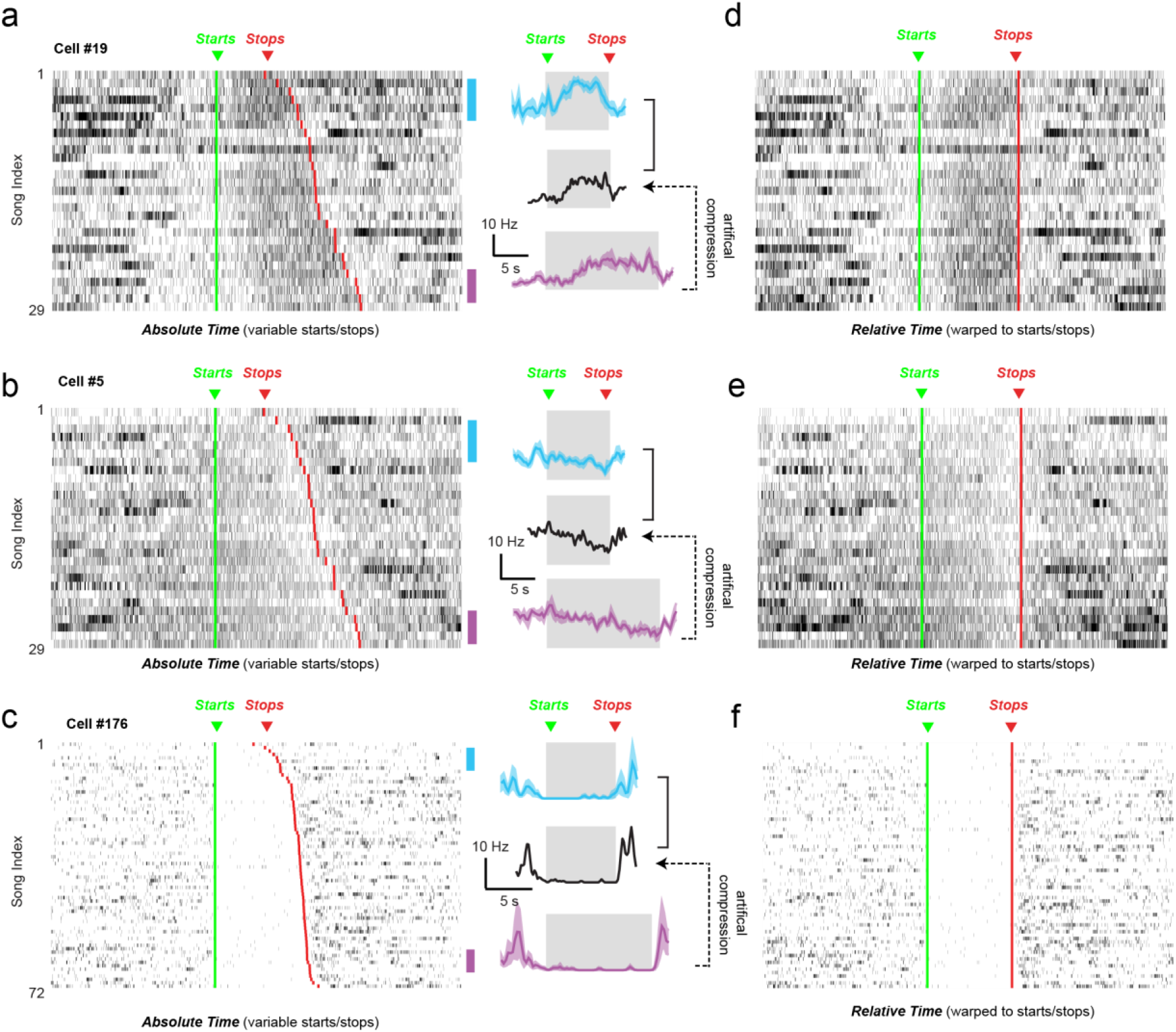
Song-modulated neurons. (**a**-**c**) Spiking raster plots for three example neurons – Cell #19 (a), Cell #5 (b), and Cell #176 (c) – across all trials. At right, a peri-song time histogram (PSTH) for song blocks representing the shortest and longest songs in the session (indicated by cyan and magenta vertical lines on right of raster plots). Black curve represents temporally compressed PSTHs from longest trials as a comparison. The magnitude of compression was chosen to match the ratio of the song durations. (**d**-**f**) Spike times of neurons in (a-c) after temporally warping to the beginning and end of song. Green and red lines indicate the onset and offset of songs, respectively.

**Extended Data Fig. 4.**
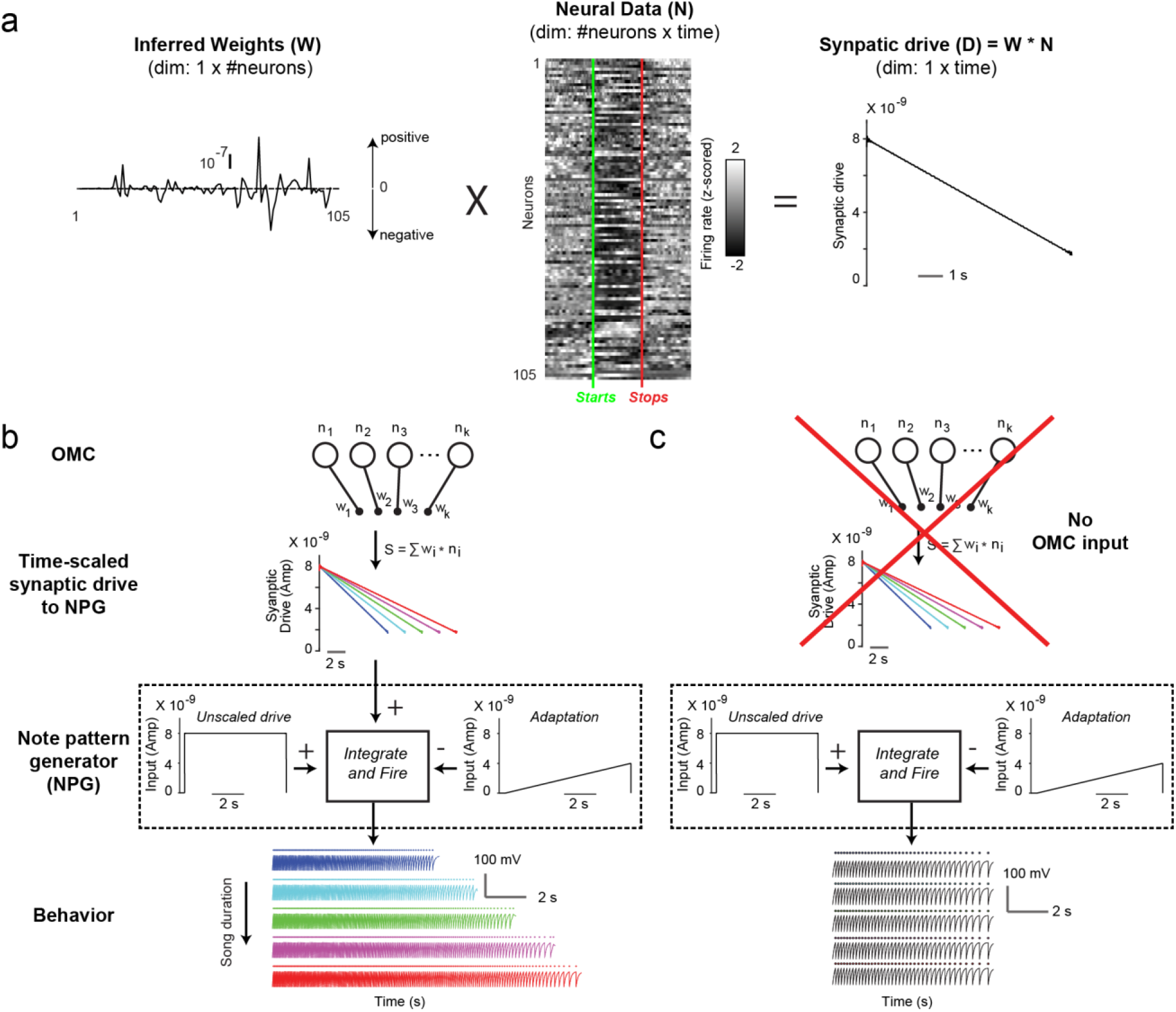
Details of the computational model. (**a**) Inferred weights (shown at left) for each song-modulated OMC neuron (shown in middle) which leads to a descending synaptic drive (shown at right) to the downstream note pattern generator. (**b**) An alternative implementation of the hierarchical model, in which the note pattern generator produces a song by combining an unscaled step-like input with a characteristic time-dependent adaptation. These inputs could be intrinsic to the pattern generator or could be inherited from a different brain area. In both cases, time-scaled OMC activity can interface with the existing note generating mechanism to produce adaptive behavioral variability. (**c**) In the absence of the OMC input, the note pattern generator can produce notes but loses flexibility resulting in songs with higher stereotypy, consistent with a partially autonomous motor control system.

